# The SAGA HAT module is tethered by its SWIRM domain and modulates activity of the SAGA DUB module

**DOI:** 10.1101/2022.12.29.522244

**Authors:** Sara T. Haile, Sanim Rahman, Benjamin C. Orsburn, Namandjé N. Bumpus, Cynthia Wolberger

**Affiliations:** Department of Biophysics and Biophysical Chemistry, The Johns Hopkins University School of Medicine, 725 N. Wolfe Street, Baltimore MD 21205; Department of Pharmacology and Molecular Sciences, The Johns Hopkins University School of Medicine, 725 N. Wolfe Street, Baltimore MD 21205

**Keywords:** transcription, chromatin, histone acetyltransferase, deubiquitinating enzyme

## Abstract

The SAGA (Spt-Ada-Gcn5 acetyltransferase) complex is a transcriptional coactivator that both acetylates and deubiquitinates histones. The histone acetyltransferase (HAT) subunit, Gcn5, is part of a subcomplex of SAGA called the HAT module. A minimal HAT module complex containing Gcn5 bound to Ada2 and Ada3 is required for full Gcn5 activity on nucleosomes. Deletion studies have suggested that the Ada2 SWIRM domain plays a role in tethering the HAT module to the remainder of SAGA. While recent cryo-EM studies have resolved the structure of the core of the SAGA complex, the HAT module subunits and molecular details of its interactions with the SAGA core could not be resolved. Here we show that the SWIRM domain is required for incorporation of the HAT module into the yeast SAGA complex, but not the ADA complex, a distinct six-protein acetyltransferase complex that includes the SAGA HAT module proteins. In the isolated Gcn5/Ada2/Ada3 HAT module, deletion of the SWIRM domain modestly increased activity but had negligible effect on nucleosome binding. Loss of the HAT module due to deletion of the SWIRM domain decreases the H2B deubiquitinating activity of SAGA, indicating a role for the HAT module in regulating SAGA DUB module activity. A model of the HAT module created with Alphafold Multimer provides insights into the structural basis for our biochemical data, as well as prior deletion studies.

## INTRODUCTION

The Spt-Ada-Gcn5 (SAGA) complex is a highly conserved eukaryotic transcriptional coactivator [1] that is recruited to promoters genome-wide to regulate expression of genes transcribed by RNA polymerase II [2–4]. SAGA activates transcription through its two enzymatic activities, histone acetylation and deubiquitination, as well as by recruiting TATA-binding protein (TBP) to promoters [5–7]. The SAGA complex comprises 19 subunits, which are organized into functional modules, and has a total molecular weight of about 1.8 MDa. The histone acetyltransferase (HAT) module, which acetylates multiple lysines on histone H3 tails, contains the catalytic Gcn5 subunit bound to three additional proteins: Ada2, Ada3, and Sgf29 [8, 9]. The four-protein deubiquitinating (DUB) module removes monoubiquitin from histone H2B K123(yeast)/K120(human) through activity of its ubiquitin protease subunit, Ubp8 [5]. In addition to the catalytic modules, SAGA contains a core module that recruits TBP and a large protein, Tra1, which is also part of other transcriptional coactivator complexes [10, 11].

Recent structures of the SAGA complex from yeast [12, 13] and humans [14] have provided a wealth of insights into the architecture of the SAGA core. However, the densities for the HAT and DUB modules were poorly resolved, indicating that they are quite mobile relative to the remainder of SAGA. Crystal structures of the DUB module [15–17] could be docked into maps to produce a model of its relation to the SAGA core. The SAGA structures also revealed how the yeast DUB module subunit, Sgf73, tethers this subcomplex to the remainder of SAGA via a central stretch of about 100 residues that is buried within the SAGA core [13]. Comparable modeling of the HAT module and its relation to the SAGA core, however, was not possible. Structural information to date on the HAT module, is limited to crystal structures of the Gcn5 catalytic domain, as well as a subset of individual domains of other HAT module subunits (reviewed in [18]). Moreover, unlike the DUB module, none of the HAT module subunits is integral to the SAGA core. One study tentatively assigned unaccounted-for helical density, which was located adjacent to core subunit Taf6, to Ada3 [12]. However, the lack of resolved side chains in the map makes this identification uncertain.

In addition to the acetyltransferase activity of the catalytic subunit, Gcn5 [19], the four subunits of the SAGA HAT module contain multiple chromatin-interacting domains. Gcn5 contains a bromodomain, which binds to acetyl lysine and regulates the enzyme’s specificity for particular lysine residues in histone H3 [20, 21]. The other HAT module subunits, Ada2, Ada3 and Sgf29 [22–25], modulate the activity and specificity of Gcn5. On its own, Gcn5 acetylates histone H3 tail peptides but is only active on nucleosomes when it is in complex with both Ada2 and Ada3 [8, 26]. While Sgf29 is not required for acetylation *in vitro*, it contains a tandem Tudor domain that recognizes trimethylated histone H3K4 and helps maintain levels of H3 acetylation *in vivo* [24, 27, 28], and promotes processive, multisite acetylation by Gcn5 i*n vitro* [29]. Ada2 contains three chromatin reader domains: the ZZ (zinc finger), SANT and SWIRM domains. The Ada2 SANT domain contributes to full HAT module activity by enhancing binding of Gcn5 to its co-substrate, acetyl-CoA [30–32], but the function of the ZZ domain remains unknown. A structure of the yeast Gcn5 core bound to the Ada2 SANT and ZZ domain shed light on the possible mechanism by which the SANT domain contributes to Gcn5 activity [32], but the function of the adjacent ZZ domain remains unknown.

The SWIRM (Swi3, Rsc8 and Moira) domain was first identified through a computational sequence profile analysis as a chromatin-interacting domain found in multiple proteins involved in regulating transcription [33]. The ~80 residue human ADA2 SWIRM domain adopts a histone-like fold and was shown to bind mono- and di-nucleosomes [34]. The SWIRM domains found in the LSD1 histone demethylase and the Swi3 subunit of the SWI/SNF nucleosome remodeling enzyme binds histone tail peptides and mononucleosomes [35, 36]. However, deletion and pull-down studies aimed at elucidating the architectural network of the SAGA complex suggested a different potential role for Ada2 and its SWIRM domain. A mass spectrometry-cross linking study [9] showed that deleting Ada2 in yeast abrogated association of the other three HAT module subunits with the SAGA complex, while deleting Gcn5 or Sgf29 did not, pointing to a specific role for Ada2 in anchoring the HAT module to the remainder of the coactivator complex. In a separate study [37], deleting the Ada2 SWIRM domain did not disrupt Ada2 association with other HAT module subunits, but resulted in an almost complete loss of association with core SAGA proteins, Sgf73 and Taf12.

While the two catalytic activities of SAGA are distinct, there is some evidence of crosstalk between the HAT and DUB modules. In humans, depletion of GCN5 results in loss of USP22, the human ortholog of Ubp8, from the SAGA complex [38]. A polyglutamine expansion in human DUB module subunit ATXN7, which gives rise to the neurodegenerative disease Spinocerebellar ataxia type 7 (SCA7), causes a significant decrease in SAGA acetyltransferase activity [39]. A deletion in the yeast ortholog, Sgf73, that causes the DUB module to dissociate from SAGA results in a twofold decrease in SAGA HAT activity [37]. While these observations point to a role for the DUB module in modulating SAGA HAT module activity, the impact of the HAT module on DUB module activity has not been explored.

Here we characterize the role of the Ada2 SWIRM domain in both tethering the HAT module to SAGA complex and in enzymatic activity. Using mass spectrometry, we show that deleting the Ada2 SWIRM domain leads to disassociation of the HAT module from the rest of the SAGA complex, although a very small proportion of SAGA complexes retain the HAT module even in the absence of the SWIRM domain. By contrast, the Ada2 SWIRM domain does not appear to play a role in the integrity of the related ADA complex, which contains the SAGA HAT module subunits in association with two additional proteins, Ahc1 and Ahc2. Loss of the HAT module due to deletion of the SWIRM domain decreases the H2B deubiquitinating activity of SAGA, indicating a role for the HAT module in modulating SAGA DUB module activity. A model of the HAT module generated using AlphaFold Multimer provides insights into the structural basis for the biochemical data. Our study provides a model for understanding how intracomplex regulation occurs within SAGA.

## METHODS

### Mutagenesis

#### Cloning of ΔSWIRM HAT module

The polycistronic vector pST44 containing *Saccharomyces cerevisiae* Ada2, Ada3 and Gcn5 was obtained from Song Tan [40]. The ΔSWIRM HAT module was generated using around the horn PCR mutagenesis with primers that excluded the SWIRM domain (amino acids 349-434).

#### Preparation of yeast strains

The FLAG-tagged WT and ΔSWIRM SAGA-containing yeast strains, yHY81 and yHY83 respectively Table 2.2), used for the pull downs were obtained from the laboratory of Dr. Steven Hahn [37]. All other experiments done with SAGA used strains derived from BY4741 (Open BioSystems). A TAP tag along with a HIS3MX6 selection marker were added to the C terminus of the Spt7 subunit via homologous recombination. SAGA ΔSWIRM was prepared by deleting the SWIRM domain via homologous recombination and adding a KanMX selection marker. Strains were verified using PCR.

### Protein expression and purification

#### Purification of endogenous SAGA

All SAGA preparations (WT, ΔSWIRM, with and without active DUB module) were purified from endogenous expression levels in *S. cerevisiae* via the Spt7-TAP tag. Yeast were grown in 6 L YPD and harvested at an OD_600_ 4-6. Pellets were resuspended in 40 mM HEPES pH 7.5, 300 mM NaCl, 100 mM NaF, 0.6 mM Na_3_VO_4_ and flash frozen in liquid nitrogen. Frozen yeast pellets were lysed in a cryogenic freezer mill (SPEX Sample Prep 6875D). Lysates were thawed, mixed with equal volume lysis buffer (50 mM HEPES pH 7.5, 300 mM NaCl, 20% Glycerol, 0.2% Tween 20, 50 mM NaF, 0.6 mM Na_3_VO_4_, 0.25 mM β-mercaptoethanol, 1 mM 4-(2-Aminoethyl)-benzenesulfonylfluoride hydrochloride (AEBSF) and 1 complete ULTRA protease inhibitor tablet (Sigma-Aldrich)) then clarified by centrifugation at 14,000 *x g* for 30 minutes. SAGA was purified as described [13] with several modifications. Briefly, clarified lysates were incubated with 2 mL of IgG Sepharose 6 Fast Flow (Cytiva) for 3 hours then washed with 200 mL of IgG wash buffer (40 mM HEPES pH 7.5, 400 mM NaCl, 10% Glycerol, 0.2% NP40, 0.25 mM β-mercaptoethanol) followed by addition 30 mL of TEV cleavage buffer (40 mM HEPES pH 7.5, 300 mM NaCl, 20% Glycerol, 0.5 mM BME). To cleave the Spt7 TAP tag and release SAGA from the beads, 1.5 mg of TEV protease was added and slurry was brought to 15 mL with TEV cleavage buffer and incubated at 16°C for 1.5 hours. The TEV cleaved IgG eluate was loaded onto a 5 mL HiTrap Q column (Cytiva) and eluted using a 150 mM to 1 M NaCl gradient over 20 column volumes. Peak fractions were concentrated using centrifugal concentrators and aliquots were flash frozen in liquid nitrogen and stored at −80°C.

#### Gcn5/Ada2/Ada3 HAT module purification

A polycistronic plasmid containing Gcn5, Ada2 and Ada3 was obtained from the laboratory of Dr. Song Tan. The heterotrimeric HAT module was purified as described [40] with some changes. The purification involves overexpression in *E. coli* followed by affinity purification via a hexahistidine tag on Ada3 and an anion exchange step. The following changes were applied to the published protocol: plasmids were transformed in R2 pLysS cells, cells were lysed with a MicroFluidizer^®^ (Microfluidics) and soluble lysates were batch purified using HisPur^™^ Ni-NTA resin (ThermoFisher) followed by a Source Q column (Cytiva). Purified HAT module was aliquoted, flash-frozen in liquid nitrogen and stored at −80°C.

#### Nucleosome assembly

Nucleosomes were assembled with either 147 base pair 601 Widom DNA sequence [41]. DNA was prepared by phenol extraction^5^. Nucleosomes were prepared by reconstituting *X. Leavis* histones into octamers followed by reconstitution into nucleosomes via a salt gradient dialysis with 601 Widom DNA as described [42].

### Electrophoretic mobility shift assays

Binding between nucleosomes and isolated HAT module was performed by incubating 0-4 *μ*M HAT with 50 nM nucleosome at 30°C for 30min in binding buffer (20 mM HEPES pH 7.5, 150 mM potassium Acetate, 4 mM MgCl_2_, 0.2 mg/mL bovine serum albumin (BSA), 1 mM Dithiothreitol (DTT). The samples were loaded onto a 4% TBE gel that had been pre-run for 60 min at 4°C in 0.5 X TBE and then run for 90 min following sample loading. Complexes were visualized using SYBR Gold, which stains DNA. Experiments were done in triplicate and gel images were recorded with a ChemDoc imager (BioRad) and analyzed using Image Lab 6.1 by quantitating the unbound nucleosome band.

### Acetylation kinetics

Steady-state kinetics of acetyltransferase activity on nucleosomes were measured using a radioactive filter binding assay [43]. Varying concentrations of nucleosome and 50 nM HAT module were incubated at 30°C for 10 minutes in buffer containing 100 mM HEPES pH 7.5, 300 mM potassium acetate and 1 mM DTT followed by addition of 10 *μ*M acetyl-CoA in a 1:10 ratio of ^3^H radiolabeled acetyl-CoA (PerkinElmer) to cold acetyl-CoA. 25 *μ*L reactions were blotted onto P81 filter paper at indicated time points (generally, 1, 30 and 60 min). For end point assays, samples were blotted after 3 hours. Filter papers were washed with 50 mM sodium bicarbonate using a vacuum wash apparatus and dried under vacuum. Samples were then added to vials containing scintillation fluid and counted. Counts were converted to concentration of acetylated nucleosome using a standard curve of known concentration of acetyl-CoA. Experiments were performed in duplicate, normalized to enzyme concentration, and plotted as a function of substrate concentration in GraphPad Prism 9.

Steady-state kinetic measurements using H3 tail peptides were conducted using a continuous spectrophotometric assay [29, 43]. 50 *μ*L reactions were prepared with buffer containing 50 mM HEPES pH 7.6, 50 mM potassium acetate, 5 mM MgCl_2_, 0.2 mM thiamine pyrophosphate, 1 mM DTT, 0.2 mg/mL BSA, 0.2 mM NAD^+^, 2.5 mM pyruvate, 1 *μ*L of 0.57 U/mg pyruvate dehydrogenase (Sigma) at 12.5 mg/mL. 50 nM HAT module was incubated in buffer along with 0-700 mM H3 peptide (amino acids 1-21) (AnaSpec) at 30°C in 384-well plates (Greiner Bio-One) for 10min. Reactions were started by adding acetyl-CoA to a final concentration of 200 *μ*M. Absorbance at 340 nm was continuously monitored by a POLARstar Omega plate reader (BMG Labtech) for 60 minutes. Absorbance at 340 nm was converted to molar concentration of NADH using a standard curve. Rates were measured in triplicate, normalized to enzyme concentration, and plotted as a function of substrate concentration in GraphPad Prism 9.

### Deubiquitinating Assay

DUB activity of wild type and Ada2 ΔSWIRM HAT module was measured as described [17] with the following modifications: sodium acetate was used instead of NaCl and the enzyme concentration was 100 nM. Experiments done with excess ubiquitinated nucleosome used purified recombinant mononucleosomes containing histone H2B ubiquitinated at K120 (H2B-K120Ub) (EpiCypher, 16-0370) at 400 nM and 25 nM SAGA. Single turnover experiments performed with limiting ubiquitinated nucleosome used purified HeLa mononucleosomes (EpiCypher 16-0002) with 100 nM SAGA. Purified HeLa nucleosomes were used at 5*μ*M; western blot was used to determine 1% of the population contained H2B-Ub so the final approximate H2B-Ub nucleosome concentration was ~50 nM. Samples were visualized by western blot with an anti H2B-K120Ub antibody (Cell Signaling #5546).

### Electrophoretic Mobility Shift Assays (EMSA)

Binding reactions containing 0-4 *μ*M recombinant HAT module and 50 nM nucleosome core particle at 30°C for 30 min in binding buffer containing 20 mM HEPES pH 7.5, 150 mM potassium acetate, 4 mM MgCl_2_, 0.2 mg/mL BSA, and 1 mM DTT. For visualization the samples were loaded onto a 4% TBE gel that pre ran for 60 min at 4°C in 0.5 X TBE then run for 90 min with the samples. Gels were stained with SYBR Gold (Invitrogen). Experiments were done in triplicate and gel images were analyzed using Image Lab 6.1 by quantitating the unbound nucleosome band.

### Mass spectrometry

#### Mass spectrometry on purified SAGA

An amount of 1 *μ*g of each sample was brought to 50 *μ*l in 20 mM ammonium bicarbonate pH 8, reduced with 5 *μ*l of 50 mM DTT for 1 hour at 60°C, and alkylated with 5 *μ*l of 50 mM iodoacetamide (IAA). Each sample was then digested with 0.25 *μ*g of Trypsin/LysC overnight at 37°C. Samples were then acidified and buffer-exchanged over an Oasis micro HLB solid phase extraction plate to remove salts and contaminants, then dried down in a SpeedVac (ThermoFisher Scientific). Samples were reconstituted in 10 *μ*L of 2% acetonitrile, 0.1% formic acid and a volume of 2 *μ*L (approximately 200 ng on column) and analyzed by reverse-phase chromatography tandem mass spectrometry on an Easy-nLC 1000 UPLC interfaced with a Orbitrap-Fusion Lumos mass spectrometer (Thermo Fisher Scientific). Peptides were separated using a 2%–90% acetonitrile in 0.1% formic acid gradient over 120 min at 300 nl/min. The 75 um x 15 cm column (PicoFrit Self pack emitter, New Objective) was in house packed with ReproSIL-Pur-120-C18-AQ (3 *μ*m, 120 Å bulk phase, Dr. Maisch). Survey scans of precursor ions were acquired from 350-1400 m/z at 120,000 resolution at 200 m/z. Precursor ions were individually isolated within 0.7 m/z by data dependent monitoring with a 15s dynamic exclusion, and fragmented using an HCD activation collision energy 30. Fragment ions were analyzed at 30,000 resolution.

Data were processed by Proteome Discoverer v2.4 (PD2.4, ThermoFisher Scientific) and searched with Mascot v.2.6.2 (Matrix Science, London, UK) against the SwissProt 2021 Yeast database. Search criteria included trypsin enzyme, one missed cleavage, 5 ppm precursor mass tolerance, 0.02 Da fragment mass tolerance, with carbamidomethylation on C as fixed and oxidation on M, deamidation on N or Q as variable modifications. Peptide identifications from the Mascot searches were processed and imported into Scaffold (Proteome Software Inc.), validated by ProteinProphet (Keller, A et al Anal. Chem. 2002;74(20):5383-92) to filter at a 95% confidence on peptides and proteins

#### FLAG pull down followed by mass spectrometry

Yeast containing FLAG-tagged Ada2 WT or Ada2 ΔSWIRM were a gift from Steven Hahn. FLAG-Ada2 pull downs were performed as described [37] with some modifications^2^. Briefly, yeast were grown in SD -Ura media (Sunrise Science) to an OD_600_ of 1-2 and lysed in a cryogenic freezer mill as described in the previous section. An amount of 2-4 mg of whole cell extract as incubated with 25 *μ*L ANTI-FLAG^®^ M2 affinity gel (Millipore Sigma) overnight at 4°C. FLAG resin was washed with Trisbuffered saline (TBS) and eluted with 25 *μ*L or 2 mg/mL 3X FLAG peptide (Fisher Scientific). Samples were analyzed by mass spectrometry as described below.

Two technical replicates were performed using the FLAG elution. Approximately 50 micrograms of protein were prepared for liquid chromatography – mass spectrometry (LCMS) analysis using the S-Trap Mini kit (ProtiFi, Long Island New York). Briefly, the protein was quantified using a BCA microplate assay (Pierce) to estimate total protein abundance. The mixture was adjusted to a volume of 50 *μ*L using the concentrated STrap lysis buffer with a total concentration of approximately 5% SDS. The solution was acidified with phosphoric acid and diluted in 90% LCMS grade methanol/water to a total volume of 400 *μ*L according to vendor instructions. The proteins were adhered to the STrap quartz fiber by centrifugation followed by two washes with the 90% methanol solution. A solution containing 5 *μ*g of sequencing grade trypsin (Promega) in 100 mM triethylammonium bicarbonate (TEAB) was briefly centrifuged into the S-Trap and the trap was incubated at 47°C for 1 hour for digestion. Following this step, the trap was loaded with 50 *μ*L of elution buffer 1 (50 mM TEAB, pH 8.4) and centrifuged to release digested peptides. The final step was repeated twice with elution buffers 2 (0.2% formic acid in LCMS grade water) and 3 (50% LCMS grade acetonitrile/water mixture with 0.1% formic acid), respectively. The peptide mixture was lyophilized to dryness with vacuum centrifugation (Eppendorf). Approximately 200 ng of peptide from each sample was analyzed by nanoflow liquid chromatography coupled to a Bruker TIMSTOF Flex mass analyzer operating in data dependent acquisition mode using a 1.1 second cycle time. The peptides were desalted online using a custom PepSep 2cm 100 micron Reprosil C-18 column and separated on an IonOpticks Aurora 25cm C-18 column using a total gradient length of 2 hours at 200 nanoliters/minute. The resulting .d files were processed with FragPipe 17.1 using the default closed search settings for TIMSTOF DDA analysis in MSFragger. Quantitative data was obtained through the use of IonQuant and employed both Match Between Runs and MaxLFQ using the default parameters.

The third replicate was performed by cleaving the remaining protein bound to the FLAG resin and analyzed by mass spectrometry as follows. Each sample was brought up to 40 *μ*L in 10 mM TEAB pH 8. The samples were then reduced with 5 *μ*L of 50 mM DTT for 1 hour at 60°C and alkylated with 5 *μ*L of 50 mM iodoacetamide. 2 *μ*L of the SP3 beads were then added to each sample and the proteins were allowed to bind as described [44] assuming up to 10 *μ*g of material. Digestion buffer (20 *μ*g Trypsin/2 mL 10 mM TEAB) was added, and the samples were allowed to digest overnight at 37°C. The beads were bound to the magnet and the supernatant containing the peptides were extracted and dried in a SpeedVac (ThermoFisher Scientific). Samples were brought up in 50 *μ*L of 100 mM TEAB and labeled with TMT10plex (LOT VG306772, ThermoFisher Scientific) with half the label amount as in the attached protocol (a full recipe labels up to 100 *μ*g of material). Briefly, labels 127N and 131 were brought up in 41 *μ*L of anhydrous acetonitrile and 20 *μ*L of 127N was added to sample Ada2 WT and 20 *μ*L of 131 was added to sample Ada2 ΔSWIRM. The reaction was allowed to proceed for 1 hour before being quenched with 4 *μ*L of 5% hydroxylamine each for 15 minutes, after which the two samples were mixed. The samples were dried in a SpeedVac and then rehydrated in 100 *μ*L of 10 mM TEAB. This sample was subjected to stepped fractionation on an Oasis HLB micro elution plate (Waters, Milford MA) where fractions at 10, 25, and 75% acetonitrile in 10 mM TEAB were successively eluted from the Oasis plate. These three fractions were dried and brought up in 15 *μ*L of 2% acetonitrile, 0.1% formic acid and placed in the autosampler of the Easy-nLC 1000. The three fractions were analyzed by reverse-phase chromatography tandem mass spectrometry on an Easy-nLC 1000 UPLC interfaced with a QE-Plus Orbitrap mass spectrometer (Thermo Fisher Scientific) as described above, however, using a 90 min gradient with 70K resolution for MS and 35K resolution for MS2, with a top 15 precursor setting on a QEPlus mass spectrometer (ThermoFisher Scientific). The combined data from the 3 fractions was searched using Proteome Discoverer version 2.5 along with Mascot version 2.8 with the SwissProt Yeast database allowing variable modifications for Oxidation on M and Deamidation on N and Q as well as TMT 6 Plex on K and with fixed Carbamidomethylation on C and TMT 6 Plex for N-terminal peptides. One missed cleavage was allowed at K or R and data was filtered at the 1% FDR level. The ratios for proteins in samples WT over ΔSWIRM were calculated using data normalized for total peptide amount as the standard setting in the software.

### Structure prediction of the isolated HAT Module

The HAT module structure was predicted with AlphaFold-Multimer [45], using a local installation of ColabFold [46]. The full sequences of yeast Gcn5, Ada2, Ada3, and Sgf29 were used and a 1:1:1:1 stoichiometry assumed. Structure predictions of the HAT module were carried out using templates from the Protein Data Bank with six recycling stages, after which the top five structures were subjected to energy minimization using the Amber99sb force field [47]. Visual inspection and pLDDT scores show minimal differences between the top five structures (SI Figure 1).

## RESULTS

### The Ada2 *SWIRM domain tethers the HAT module to the yeast SAGA complex*

A previous study [37] had shown that SAGA subunits, Taf12 and Sgf73, only co-precipitated with Ada2 when its C-terminal SWIRM domain was present, suggesting a role for this SWIRM domain in connecting the HAT module to the TFIID-like core of SAGA [37]. To better quantify the role of the Ada2 SWIRM domain in tethering the HAT module to the remainder of the SAGA complex, we used mass spectrometry to quantitate all SAGA complex subunits. The SAGA complex was affinity purified from yeast strains containing a TAP tag on SAGA core subunit Spt7 and then analyzed by mass spectrometry. The relative proportion of each subunit was compared for SAGA complex purified from strains containing WT Ada2 versus a strain containing a deletion of the Ada2 SWIRM domain (ΔSWIRM). Mass spectra on SAGA purified from ΔSWIRM yeast showed all HAT module components to be depleted by about seven-fold compared to wild type (Figure 1A). All other SAGA proteins were present at wild type levels in the ΔSWIRM SAGA complex. This depletion of just the HAT module proteins from the SAGA complex upon deletion of the Ada2 SWIRM domain indicates that the SWIRM domain plays a central role in tethering all four HAT module subunits to the rest of the SAGA complex.

**Figure 1:**
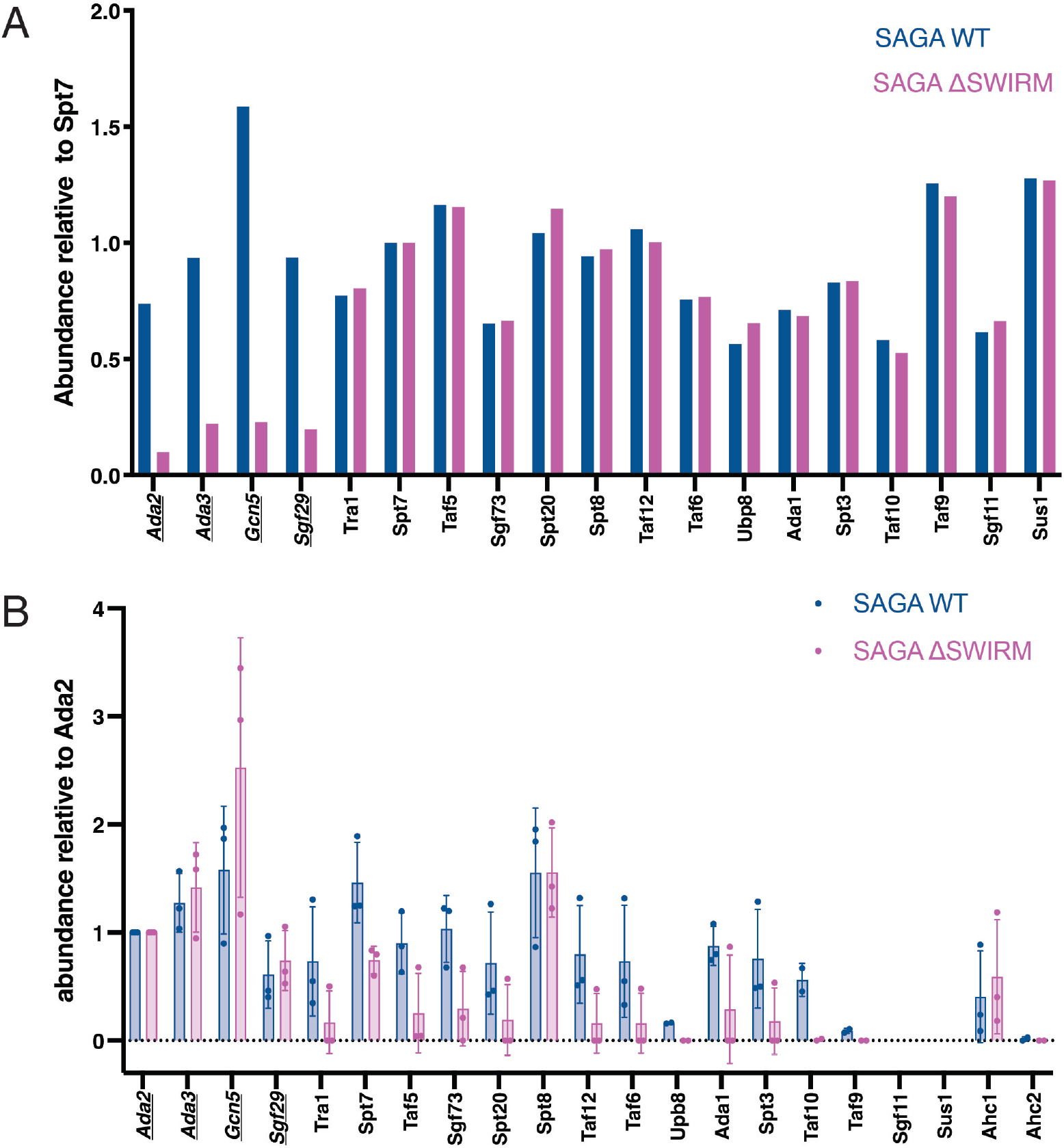
SWIRM domain is responsible for tethering the HAT module to the SAGA complex. (A) LC-MS/MS on purified WT and ΔSWIRM SAGA complexes, with total spectrum counts normalized to Spt7. HAT module proteins are underlined. (B) LC-MS/MS of Ada2-FLAG pulled downs performed in triplicate with two different types of analysis. In one experiment, FLAG-tagged Ada2 and associated proteins were Trypsin digested from FLAG resin and analyzing by tandem mass tag (TMT) labelling and a QEPlus mass spectrometer. In a second experiment, the complex was eluted from beads with the FLAG peptide and analyzed by a TIMSTOF Flex mass spectrometer. Data represent abundances collected by TMT labelling (1 replicate) and by TIMSTOF (2 replicates).

In addition to being a part of the SAGA complex, Gcn5, Ada2, Ada3, and Sgf29 can also associate with proteins Ahc1 and Ahc2 to form a smaller complex known as ADA [9, 48]. It is not known what regulates incorporation of the HAT module subunits into the ADA versus the SAGA complex. To determine if the Ada2 SWIRM domain also plays a role in regulating HAT module association with Ahc1 or Ahc2, we used yeast strains containing FLAG-tagged Ada2 WT and Ada2 ΔSWIRM domain to affinity purify the HAT module and analyzed the associated proteins using mass spectrometry. In one experiment, FLAG-tagged Ada2 and associated proteins were Trypsin digested from FLAG resin and analyzed by tandem mass tag (TMT) labelling and a QEPlus mass spectrometer. In a second experiment, the complex was eluted from beads with the FLAG peptide and analyzed by a TIMSTOF Flex mass spectrometer. Abundances in both approaches were normalized to Ada2 levels (Figure 1B). As expected, the other three HAT module subunits, Ada3, Gcn5 and Sgf29, were pulled down at comparable levels in the Ada2 ΔSWIRM pull down. Association of Ada2 with the remaining SAGA subunits was dramatically reduced in the Ada2 ΔSWIRM pull-down, with the notable exception of Spt8 (Figure 1B). This was unexpected, since previous crosslinking studies have not revealed any interactions between Spt8 and the HAT module [37, 49]. In recent cryo-EM structures of the SAGA complex [12, 13], Spt8 was not seen in the central module and is instead flexibly tethered to the core. Our data suggest that Spt8 may contact the HAT module in a way that is independent of the SWIRM domain.Interestingly, Gcn5 is pulled down in greater abundance by Ada2 ΔSWIRM than by the Ada2 WT. The reason for the observed difference will require further study.

The ADA complex proteins, Ahc1 and Ahc2, were also identified in the Ada2 pull downs (Figure 1B). While Ahc2 had poor coverage, likely due to its small size (15 kDa), Ahc1 was present in both the Ada2 WT and ΔSWIRM pull downs. These data indicate that the Ada2 SWIRM domain is not required for formation of the ADA complex and is therefore unlikely to contact Ahc1. We note that it was not possible to assess what proportion of the HAT module that is disassociated from the SAGA complex is bound to Ahc1, and likely Ahc2, to form the ADA complex, versus existing as free HAT module.

There was some coverage of the HAT module proteins in the SAGA-Ada2ΔSWIRM complex, although greatly reduced as compared to WT, suggesting that a subset of the SAGA complexes retained the HAT module even when the Ada2 SWIRM domain was deleted. We first compared the acetyltransferase activity of WT SAGA and SAGA-Ada2ΔSWIRM on both H3 tail peptide and recombinant nucleosome substrates. As shown in Figures 2A and 2B, SAGA-Ada2ΔSWIRM had no detectable acetyltransferase activity on either substrate as compared to background signal (Figures 2A and B). We next assayed the activity of SAGA-Ada2ΔSWIRM at a sevenfold higher concentration, to account for the approximately seven-fold decrease in SAGA complexes containing HAT module proteins as compared to wild type (Figure 1A). As shown in Figure 2C, the more concentrated preparation of SAGA-Ada2ΔSWIRM had comparable acetyltransferase activity to the more dilute WT SAGA (Figure 2C). The full recovery of acetyltransferase activity indicates that the HAT module is not completely lost upon deletion of the Ada2 SWIRM domain and suggest that the SWIRM domain does not play an important role in the HAT activity of intact SAGA. Taken together, these results suggest that the Ada2 SWIRM domain plays a key role in tethering the HAT module to rest of the SAGA complex but does not impact SAGA HAT activity.

**Figure 2:**
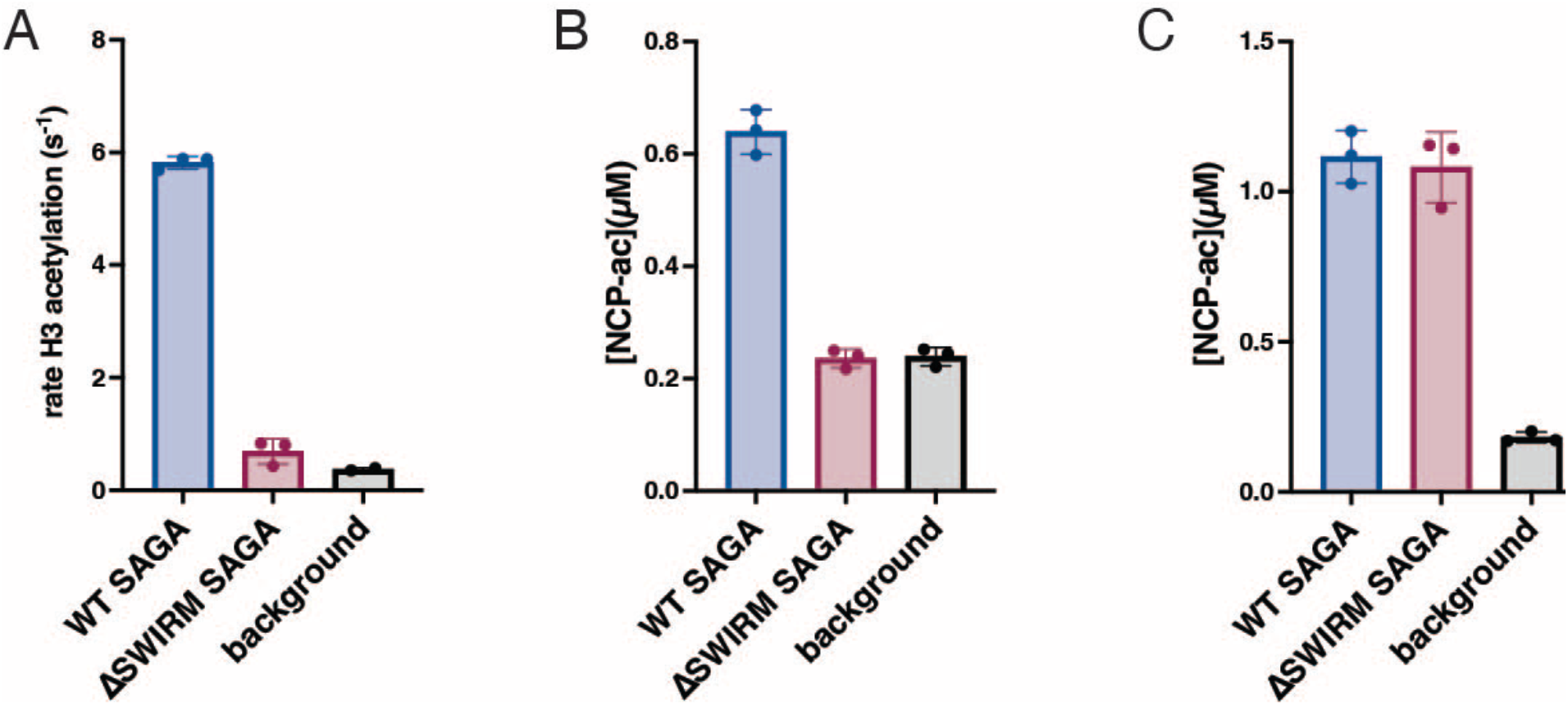
Activity lost from depletion of the HAT module can be recovered. A and B: WT and ΔSWIRM SAGA acetyltransferase activity on H3 peptides (A) and nucleosomes (B). C: SAGA acetyltransferase activity on nucleosomes where concentration of ΔSWIRM SAGA is increased seven-fold.

### The SWIRM domain only modestly affects HAT module’s ability to bind and acetylate nucleosomes

The LSD1 SWIRM domain binds histone tails and the Swi3 SWIRM domain binds mononucleosomes [35, 36]. In light of these studies pointing to a variety of roles for SWIRM domains, we further explored whether the SWIRM domain contributes to HAT module acetyltransferase activity or ability to bind the nucleosome. We prepared the three-component *S. cerevisiae* HAT (yHAT) module containing Gcn5, Ada3, and either WT or ΔSWIRM Ada2 and compared the activity of the two complexes on nucleosomes. On a nucleosome substrate, the ΔSWIRM yHAT module had a higher turnover rate, with a *k_cat_* of 1.18 min^-1^ as compared 0.48 min^-1^ for the WT. The K_M_ of the ΔSWIRM yHAT module on nucleosomes was higher, at 57.5 *μ*M, as compared to that 32.8 *μ*M for the WT (Figure 3A and Table1). Overall, deletion of the SWIRM domain increased the catalytic efficiency of the HAT module by less than 2-fold as compared to WT (Figure 3A). We also compared the effect of Ada2 SWIRM domain deletion on the ability of yHAT to bind nucleosomes. As assayed by an electrophoretic mobility shift assay (EMSA), the ΔSWIRM yHAT complex had marginally lower affinity for recombinant nucleosome as reflected by disappearance of the free nucleosome band (compare bands at 1 and 2 *μ*M in Figures 3B and 3C). Taken together, these differences are minimal and do not suggest a role of the SWIRM domain in regulating the yHAT module’s acetyltransferase activity or ability to bind nucleosomes.

**Figure 3:**
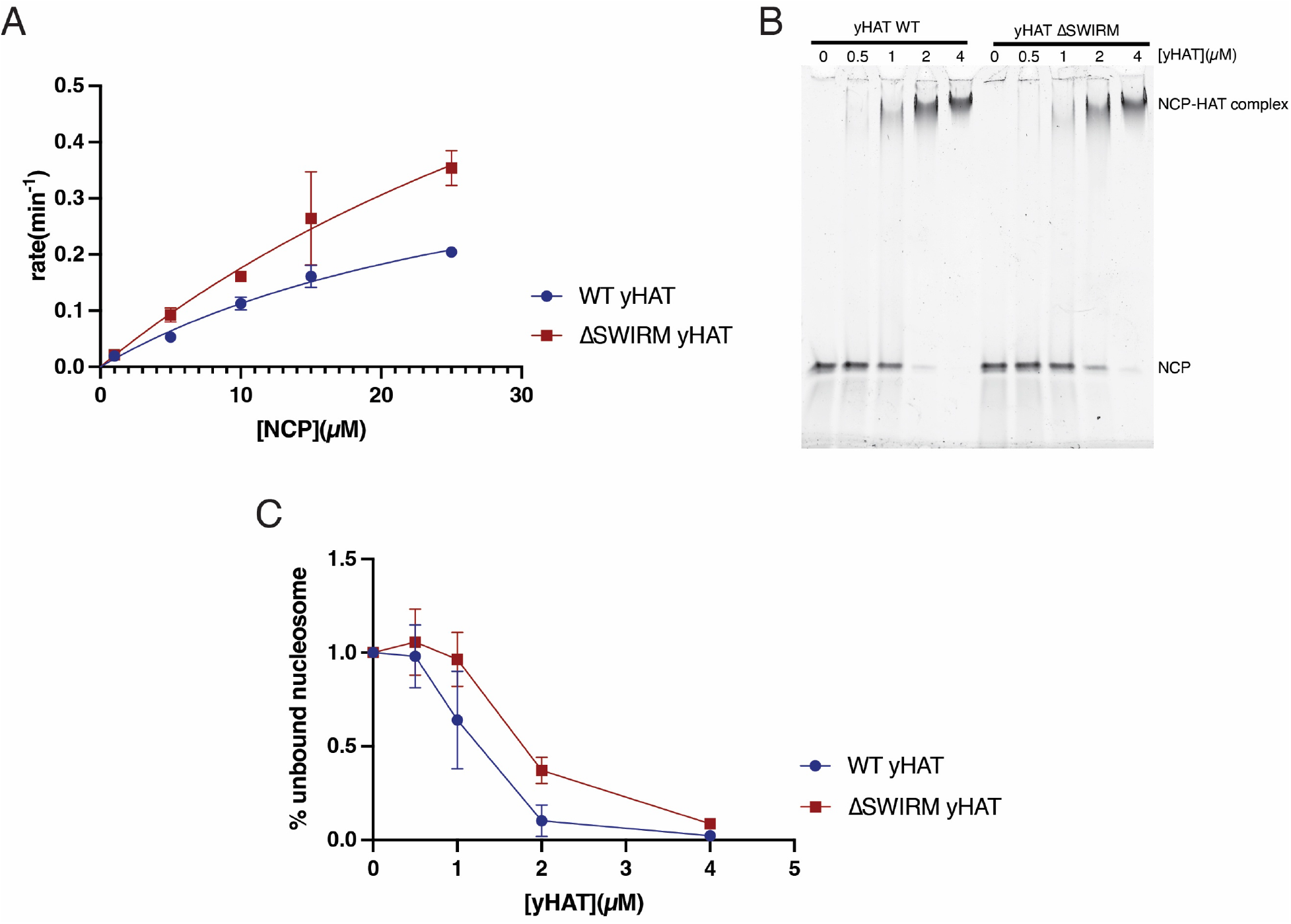
Isolated yHAT module’s ability to bind and acetylate nucleosomes is only moderately affected by deletion of the SWIRM domain. A: Acetylation of nucleosome core particles (NCP) by WT and ΔSWIRM yHAT. B: Representative electrophoretic mobility shift assay (EMSA) of binding of WT and ΔSWIRM yHAT to nucleosome core particles (NCP). C: quantitation of B with replicates.

**Table 1:**

| yHAT construct | substrate | $k_{cat}$ ($\text{min}^{-1}$) | $K_M$ ($\mu\text{M}$) | $k_{cat}/K_M$ ($\mu\text{M}^{-1}\text{min}^{-1}$) |
| --- | --- | --- | --- | --- |
| WT | NCP | $0.4816 \pm 0.1$ | $32.8 \pm 9.9$ | 0.0146 |
| ΔSWIRM | NCP | $1.18 \pm 0.66$ | $57.3 \pm 42.1$ | 0.0206 |

**Table 2.2:** List of yeast strains

| **Table 2.2: List of yeast strains** |  |  |
| --- | --- | --- |
| **Yeast strain name** | Background strain | Insertions and deletions |
| **YCW14** | BY4741 | Spt7-TAP::His3MX6 |
| **YCW20** | BY4741 | Spt7-TAP::His3MX6, Ada2 $\Delta$ residues 349-434 substituted with KanMX |
| **YHY81** | BY4705 | Sgf73-3HA::His3MX6, Δada2::KanMX, Ada2 with N terminal 3X FLAG tag |
| **YHY83** | BY4705 | SPT7-(Flag1)-TAP::TRP1, Δada2::KanMX, Ada2 ORF $\Delta$ residues 353-434 with 3xFlag on N-terminus, 491bp promoter sequence, 100bp of sequence past stop codon |

### The HAT module modulates SAGA DUB catalytic efficiency in vitro

Previous studies have demonstrated that mutations or deletions in the SAGA

DUB module affect SAGA HAT module activity [37, 39]. In yeast SAGA, HAT activity is decreased in the absence of the DUB module [37]. The impact of HAT module deletions on SAGA DUB activity, however, has not been explored. To understand how the HAT module might affect the SAGA DUB module’s ability to deubiquitinate H2B and facilitate crosstalk, we compared the deubiquitinating activity of intact WT SAGA and SAGA-Ada2ΔSWIRM, which lacks the HAT module. Recombinant nucleosomes containing monoubiquitinated histone H2B-K120 (analogous to H2B-K123 in yeast) were used to compare the DUB activity of WT and Ada2 ΔSWIRM SAGA under multiple turnover conditions. As shown in Figure 4A, there was a noticeable decrease in DUB activity in SAGA-Ada2ΔSWIRM, which is almost completely depleted of the HAT module, as compared to WT SAGA. Under the conditions tested, SAGA-Ada2ΔSWIRM deubiquitinates nucleosomes at a rate of 1.3 nM/min while the rate for WT SAGA was 4.1 nM/min, about three-fold higher.

**Figure 4:**
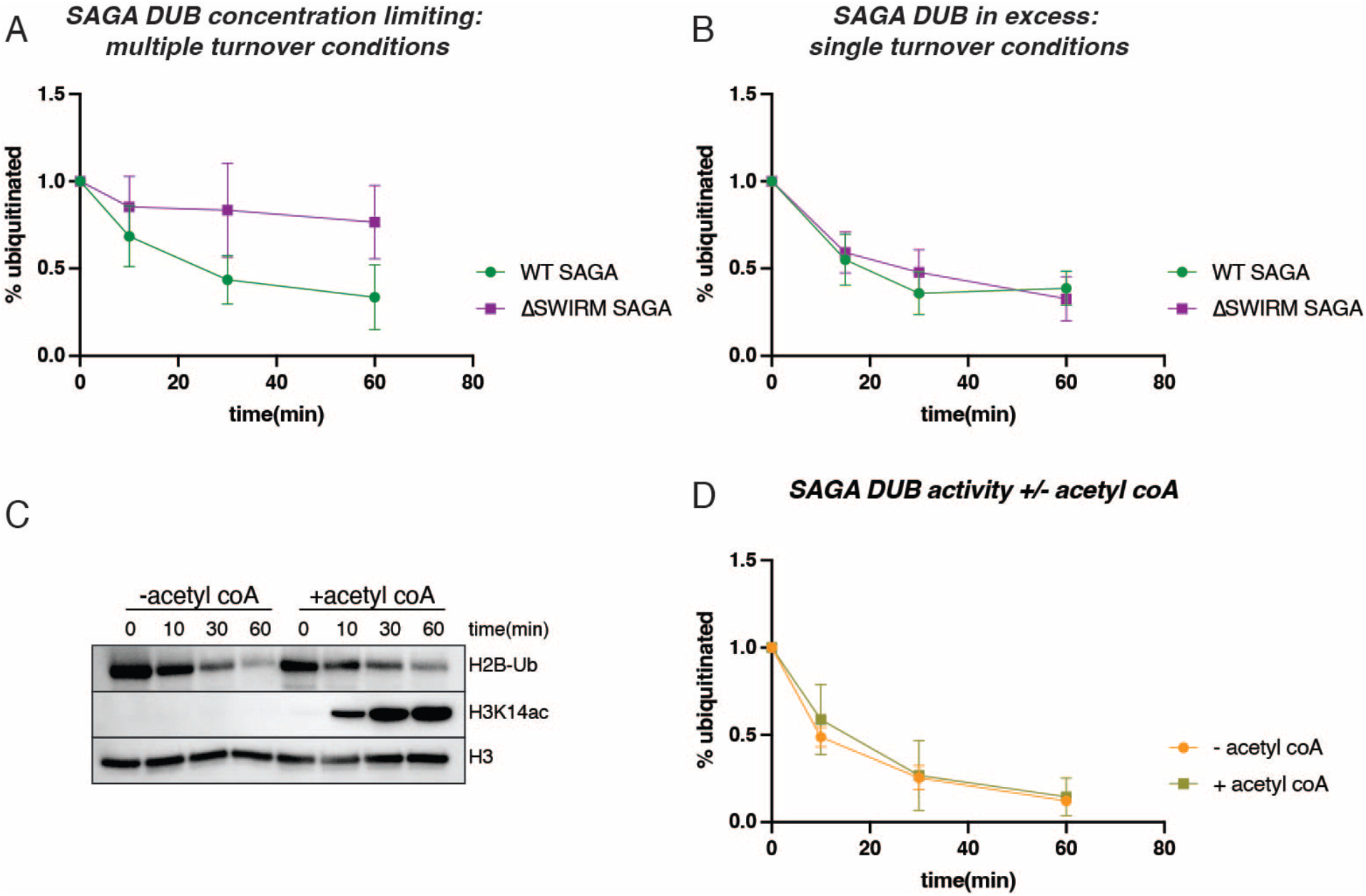
SAGA’s HAT module regulates SAGA DUB activity independent of enzymatic activity. A: Quantitation of western blots of SAGA DUB activity on recombinant H2B-Ub nucleosomes (400 nM) under conditions of limiting SAGA enzyme (25 nM) with and without the SWIRM domain. B: Quantitation of western blots of SAGA DUB activity on HeLa nucleosomes with ~50 nM H2B ubiquitinated nucleosome. SAGA DUB activity is compared with and without the SWIRM domain, with excess SAGA, 100 nM. C: WT SAGA DUB (25 nM) activity on recombinant H2B-Ub nucleosomes (400 nM) in absence and presence of acetyl CoA (10 *μ*M). D: Quantitation of C and replicate.

We also assayed SAGA DUB activity on HeLa nucleosomes under single turnover conditions, in which the concentration of nucleosome substrate was limiting and SAGA was in excess. These nucleosomes comprise a mixed population with a variety of post translational modifications, including monoubiquitinated histone H2B-K120. Under these assay conditions, SAGA DUB activity was not noticeably reduced in the absence of the HAT module (Figure 4B). This result indicates that the catalytic rate of deubiquitination is unaffected by the presence of the HAT module.

The observation that SAGA lacking the HAT module has lower H2B deubiquitinating activity under multiple turnover, but not single turnover conditions, suggests that the K_M_ for nucleosomes is higher when the HAT module is not present. While it is not known whether the SAGA HAT and DUB modules can simultaneously bind to a single nucleosome, the proximity of the density for the HAT and DUB modules in cryo-EM maps [12, 13, 50] suggests that this is a possibility. Additional interactions mediated by the SAGA HAT module with the nucleosome could thereby contribute to a lower K_M_ for the DUB module as compared with SAGA complex lacking the HAT module.

The HeLa nucleosomes used for the single turnover experiments contain a variety of post-translation modifications, including acetylated lysines on histone H3 tails. To help rule out the possibility that the presence of H3 tail acetylation impacted DUB activity in the single-turnover experiments, we assayed the DUB activity of WT SAGA on recombinant H2B-ubiquitinated nucleosomes under conditions in which the HAT module could also acetylate histone H3. WT SAGA was incubated with the nucleosomes in the presence and absence of acetyl-CoA, which is the acetyl group donor for the HAT module. Immunoblotting was used to probe for both ubiquitinated H2B and acetylated histone H3-K14 (H3-K14ac), one of the histone H3 lysines that SAGA is known to acetylate [51, 52]. As shown in Figures 4C and 4D, SAGA deubiquitinating activity is unaffected by the presence of acetylated histone H3, or by the ability of SAGA to simultaneously acetylate nucleosomes.

### A model for HAT module organization and association with SAGA

Results from the present (Figure 1) and previous studies [37] point to a key role for the Ada2 SWIRM domain in tethering the HAT module to SAGA. The location of Ada2 or the SWIRM domain within the SAGA complex is, however, unknown, since the HAT module could not be resolved in any of the four reported cryo-EM structures of SAGA [12–14, 50]. The structure of the HAT module is also not known. In order to developed a structural model for how the Ada2 SWIRM domain might anchor the HAT module to the remainder of SAGA, we used Alphafold-Multimer (AF-Multimer) [45] to generate a model of the HAT module (Figure 5A). The Predicted Local Distance Difference Test (pLDDT) [53] suggests many regions of the HAT module are predicted with moderate to high confidence (pLDDT>70) (Supplementary Figure S1A). The Predicted Alignment Error (PAE) of the highest scoring model also suggests that AF-Multimer predicted the relative positioning of ordered domains with high confidence (Supplementary Figure S1B). To validate the model, we calculated the number of previously reported interdomain crosslinks [37, 49] that were satisfied within the HAT module. Strikingly, we found that 89.1% of crosslinks reported by Han [37] were satisfied and 97.7% reported by Liu [49].

**Figure 5:**
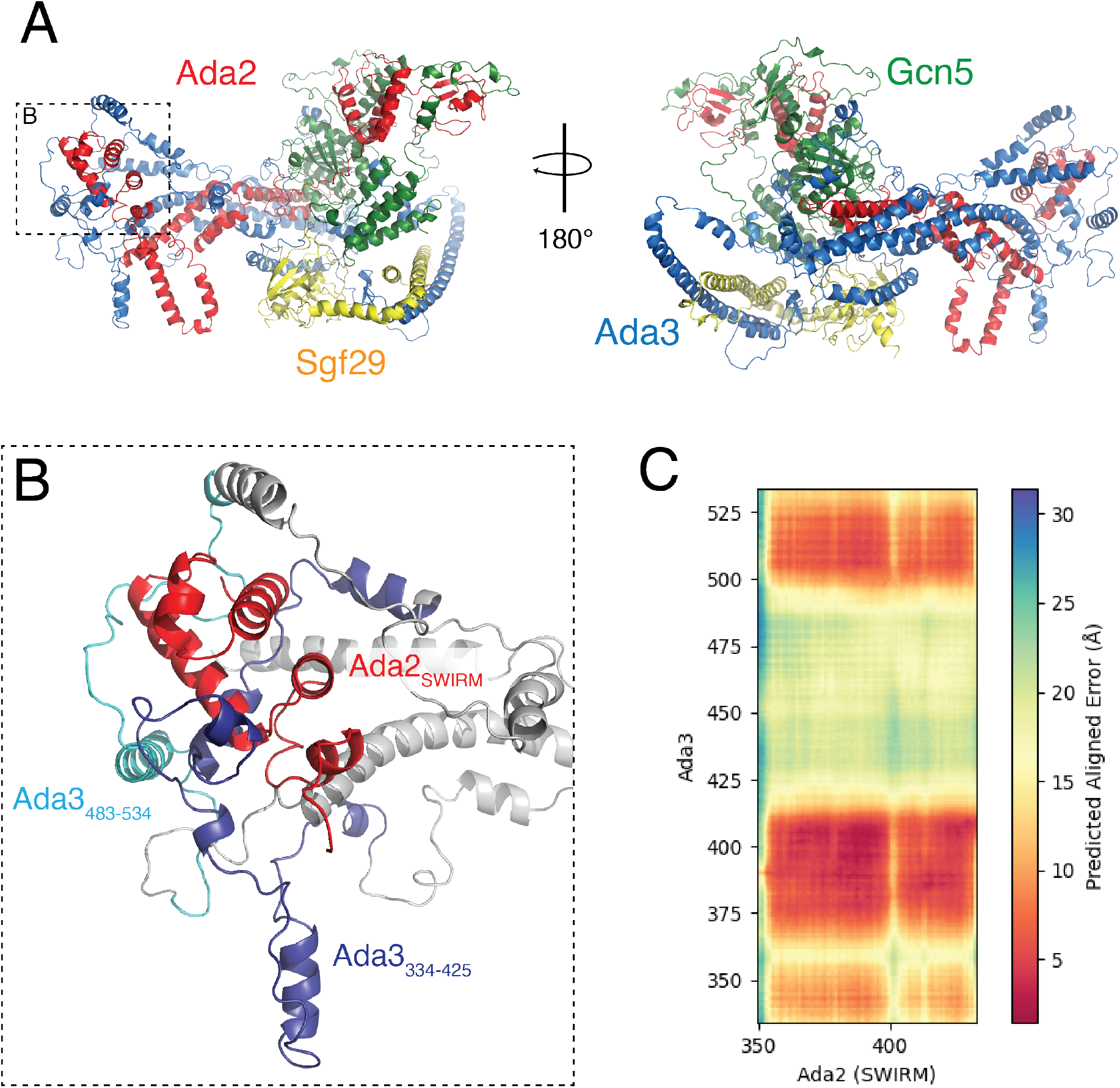
Overview of the Alphafold-Multimer structure prediction of the HAT module. A: Highest rank model of the isolated HAT module predicted from Alphafold-Multimer, containing Gcn5 (green), Ada2 (red), Ada3 (blue), and Sgf29 (gold). B: Zoom on indicated inset in panel (A) showing the positioning of Ada2’s SWIRM domain in respect to Ada3. C: Predicted Aligned Error (PAE) between Ada2’s SWIRM domain and the C-terminus of Ada3 (334-534).

The AF-Multimer model of the HAT module (Figure 5A) suggests that Gcn5, Ada2, and Sgf29 are well-structured, whereas Ada3 is predicted to contain many disordered linkers connecting helical regions that contact all HAT module components. The Ada2 SWIRM domain, whose structure is similar to the reported NMR structure (PDB ID: 2AQE) [34], forms several contacts with the Ada3 C-terminus (Figure 5B and Supplementary Figure 2A and 2B). As expected, AF-Multimer provided a high confidence model of contacts between the Gcn5 catalytic domain and the N-terminus of Ada2 (Supplementary Figure 1B) that closely resembles the crystal structure of Gcn5^72-312^ bound to Ada2^6-120^ (PDB IDs: 6CW2 & 6CW3) [32] (Supplementary Figure S2D). In contrast with the somewhat intertwined Ada2, Ada3, and Gcn5 subunits, Sgf29 and its tandem Tudor domain associate with the periphery of the complex, consistent with the observation that this subunit is not required for full activity of the HAT module [24].

The predicted model shows the Ada2 SWIRM domain (residues 349-432) to be located at one end of the HAT module complex (Figure 5B), where it interacts with portions of Ada3. Approximately 63% of the SWIRM domain is solvent-accessible (3,351 Å^2^ out of 5,288 Å^2^ of surface area) and potentially available for interactions with the core of the SAGA complex. Interestingly, the Ada2 SWIRM domain is contacted by two segments of Ada3 that, when deleted, cause the HAT module to detach from the rest of SAGA [37]. Deletion of residues Ada3 residues 334-425 or of 483-534 (Figure 5B) was shown to abrogate tethering to the SAGA core, as judged by pulldowns and immunoblotting for core subunits, Taf12 and Sgf73 [37]. By contrast, neither deletion disrupted the ability of Ada3 to co-precipitate with Ada2 or Gcn5 [37]. The proximity of these Ada3 residues to the SWIRM domain further implicates this portion of the HAT module in binding to the SAGA core.

## Discussion

Previous studies had pointed to a role for Ada2, and specifically its SWIRM domain, in anchoring the HAT module to the rest of the SAGA complex [9, 37, 54]. Here we quantitated the effects of deleting the Ada2 SWIRM domain and showed that this deletion leads to depletion of about 85% of the HAT module from the purified SAGA complex (Figure 1A). The resulting loss of acetyltransferase activity could be recovered by increasing the concentration of Ada2 ΔSWIRM SAGA by seven-fold (Figure 2). Together with the observation that deletion of the SWIRM domain did not affect nucleosome binding (Figure 3B), these results suggest the SWIRM domain primarily plays a structural role in tethering the HAT module to the remainder of the SAGA complex. This structural role is specific to SAGA, as the SWIRM domain was not essential to maintaining the integrity of the yeast ADA complex (Figure 1B), which contains the four HAT module subunits associated with the proteins, Ahc1 and Ahc2 [9].

The model of the SAGA HAT module generated with Alphafold-Multimer [45] predicts a somewhat elongated complex, with the SWIRM domain and Gcn5 located at opposite ends. This apposition would suggest that the SWIRM domain is the attachment point to the SAGA core, about which the HAT module pivots (Figure 6). Interestingly, Ada3 residues that have also been implicated in tethering the HAT module to the SAGA core [37] are also located at the distal end of the HAT module, where they contact the Ada2 SWIRM domain (Figures 5A and 5B). The proximity to the SWIRM domain of Ada3 residues that are also important for HAT module in binding to the SAGA core provides support for the validity of the predicted model. We note that, since deletion of either the Ada3 residues or the SWIRM domain has the potential to disrupt the overall structure of this region of the HAT module, it is not possible to distinguish whether the Ada2 SWIRM domain, residues within Ada3 334-425 or 483-534, or both proteins mediate direct contacts with the SAGA core.

**Figure 6:**
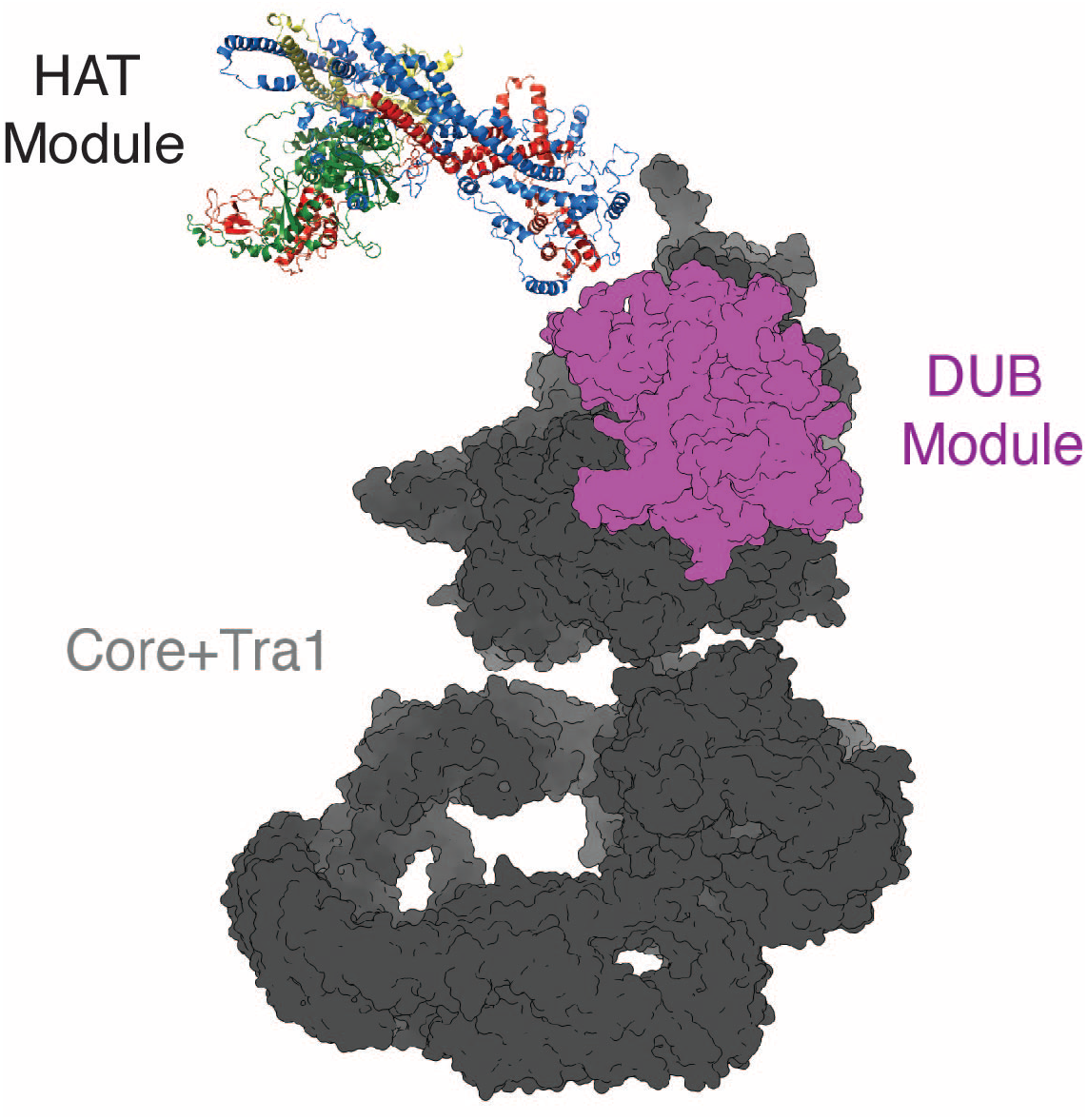
Proposed model of the SAGA HAT module positioning with respect to the SAGA core and DUB module. The Ada2 SWIRM domain and the C-terminus of Ada3 are positioned to tether the HAT module to rest of the SAGA complex. Core yeast SAGA complex from PDB ID 6T9I [18], position of DUB module from PDB ID 6TBM [12].

Our finding that the SAGA HAT module modulates SAGA DUB module activity is a directionality of crosstalk that has not previously been reported. The important role played by the Ada2 SWIRM domain in tethering the HAT module to the SAGA complex provided an opportunity to study crosstalk between the HAT module and the other catalytic subcomplex of SAGA, the DUB module. Previous studies had focused on how the DUB module affects HAT module activity. Spinocerebellar Ataxia type 7, which is a consequence of a polyglutamine expansion in the human homolog of the Sgf73 DUB module subunit, Ataxin-7, was reported to reduce the SAGA HAT module’s acetyltransferase activity [39]. In yeast, deletions in Sgf73 can either modestly stimulate or reduce HAT module activity, depending on the nature of the deletion [37]. In this study, we compared DUB module activity under different conditions in the presence and absence of the HAT module by using the ΔSWIRM SAGA, in which the HAT module is depleted. We find that the HAT module appears to module the DUB module’s catalytic efficiency by lowering the K_M_ of SAGA for H2B-ubiquitinated nucleosomes (Figures 2C and 3A). The proximity of the HAT and DUB module densities in cryo-EM structures of intact SAGA [13, 50] could explain the K_M_ effect we observe simply based the loss of additional contacts with the nucleosome. A full understanding of the interplay between the HAT and DUB modules, as well as the structural details of their attachment to the SAGA core, awaits the high-resolution structure determination of a complete SAGA complex bound to nucleosomal substrates.

## ACKNOWLEDGEMENTS

Supported by National Institutes of Health grants R35GM13093 (C.W.), 5R01AG064908 and 5R01GM103853 (N.N.B and B.C.O.), and T32GM135131 (S.R.); and a Howard Hughes Medical Institute Gilliam Fellowship (S.T.H.). The Johns Hopkins School of Medicine Mass Spectrometry and Proteomics Core is supported by National Cancer Institute center grant P30 CA006973 to the Sidney Kimmel Comprehensive Cancer Center.

## REFERENCES

[1] Y.C. Chen, S.Y.R. Dent, Conservation and diversity of the eukaryotic SAGA coactivator complex across kingdoms, Epigenetics Chromatin 14(1) (2021) 26.

[2] T. Baptista, S. Grunberg, N. Minoungou, M.J.E. Koster, H.T.M. Timmers, S. Hahn, D. Devys, L. Tora, SAGA Is a General Cofactor for RNA Polymerase II Transcription, Mol Cell 68(1) (2017) 130–143 e5.

[3] J. Bonnet, C.Y. Wang, T. Baptista, S.D. Vincent, W.C. Hsiao, M. Stierle, C.F. Kao, L. Tora, D. Devys, The SAGA coactivator complex acts on the whole transcribed genome and is required for RNA polymerase II transcription, Genes Dev 28(18) (2014) 1999–2012.

[4] L. Warfield, S. Ramachandran, T. Baptista, D. Devys, L. Tora, S. Hahn, Transcription of Nearly All Yeast RNA Polymerase II-Transcribed Genes Is Dependent on Transcription Factor TFIID, Mol Cell 68(1) (2017) 118–129 e5.

[5] K.W. Henry, A. Wyce, W.S. Lo, L.J. Duggan, N.C. Emre, C.F. Kao, L. Pillus, A. Shilatifard, M.A. Osley, S.L. Berger, Transcriptional activation via sequential histone H2B ubiquitylation and deubiquitylation, mediated by SAGA-associated Ubp8, Genes Dev 17(21) (2003) 2648–63.

[6] M.H. Kuo, J. Zhou, P. Jambeck, M.E. Churchill, C.D. Allis, Histone acetyltransferase activity of yeast Gcn5p is required for the activation of target genes in vivo, Genes Dev 12(5) (1998) 627–39.

[7] E. Larschan, F. Winston, The S. cerevisiae SAGA complex functions in vivo as a coactivator for transcriptional activation by Gal4, Genes Dev 15(15) (2001) 1946–56.

[8] R. Balasubramanian, M.G. Pray-Grant, W. Selleck, P.A. Grant, S. Tan, Role of the Ada2 and Ada3 transcriptional coactivators in histone acetylation, J Biol Chem 277(10) (2002) 7989–95.

[9] K.K. Lee, M.E. Sardiu, S.K. Swanson, J.M. Gilmore, M. Torok, P.A. Grant, L. Florens, J.L. Workman, M.P. Washburn, Combinatorial depletion analysis to assemble the network architecture of the SAGA and ADA chromatin remodeling complexes, Mol Syst Biol 7 (2011) 503.

[10] C.E. Brown, L. Howe, K. Sousa, S.C. Alley, M.J. Carrozza, S. Tan, J.L. Workman, Recruitment of HAT complexes by direct activator interactions with the ATM-related Tra1 subunit, Science 292(5525) (2001) 2333–7.

[11] M.G. Pray-Grant, D. Schieltz, S.J. McMahon, J.M. Wood, E.L. Kennedy, R.G. Cook, J.L. Workman, J.R. Yates, 3rd, P.A. Grant, The novel SLIK histone acetyltransferase complex functions in the yeast retrograde response pathway, Mol Cell Biol 22(24) (2002) 8774–86.

[12] G. Papai, A. Frechard, O. Kolesnikova, C. Crucifix, P. Schultz, A. Ben-Shem, Structure of SAGA and mechanism of TBP deposition on gene promoters, Nature 577(7792) (2020) 711–716.

[13] H. Wang, C. Dienemann, A. Stutzer, H. Urlaub, A.C.M. Cheung, P. Cramer, Structure of the transcription coactivator SAGA, Nature 577(7792) (2020) 717–720.

[14] D.A. Herbst, M.N. Esbin, R.K. Louder, C. Dugast-Darzacq, G.M. Dailey, Q. Fang, X. Darzacq, R. Tjian, E. Nogales, Structure of the human SAGA coactivator complex: The divergent architecture of human SAGA allows modular coordination of transcription activation and co-transcriptional splicing, bioRxiv (2021) 2021.02.08.430339.

[15] N.L. Samara, A.B. Datta, C.E. Berndsen, X. Zhang, T. Yao, R.E. Cohen, C. Wolberger, Structural insights into the assembly and function of the SAGA deubiquitinating module, Science 328(5981) (2010) 1025–9.

[16] A. Kohler, E. Zimmerman, M. Schneider, E. Hurt, N. Zheng, Structural basis for assembly and activation of the heterotetrameric SAGA histone H2B deubiquitinase module, Cell 141(4) (2010) 606–17.

[17] M.T. Morgan, M. Haj-Yahya, A.E. Ringel, P. Bandi, A. Brik, C. Wolberger, Structural basis for histone H2B deubiquitination by the SAGA DUB module, Science 351(6274) (2016) 725–8.

[18] J.M. Espinola-Lopez, S. Tan, The Ada2/Ada3/Gcn5/Sgf29 histone acetyltransferase module, Biochim Biophys Acta Gene Regul Mech 1864(2) (2021) 194629.

[19] J.E. Brownell, J. Zhou, T. Ranalli, R. Kobayashi, D.G. Edmondson, S.Y. Roth, C.D. Allis, Tetrahymena histone acetyltransferase A: a homolog to yeast Gcn5p linking histone acetylation to gene activation, Cell 84(6) (1996) 843–51.

[20] D.E. Sterner, P.A. Grant, S.M. Roberts, L.J. Duggan, R. Belotserkovskaya, L.A. Pacella, F. Winston, J.L. Workman, S.L. Berger, Functional organization of the yeast SAGA complex: distinct components involved in structural integrity, nucleosome acetylation, and TATA-binding protein interaction, Mol Cell Biol 19(1) (1999) 86–98.

[21] A.M. Cieniewicz, L. Moreland, A.E. Ringel, S.G. Mackintosh, A. Raman, T.M. Gilbert, C. Wolberger, A.J. Tackett, S.D. Taverna, The bromodomain of Gcn5 regulates site specificity of lysine acetylation on histone H3, Mol Cell Proteomics 13(11) (2014) 2896–910.

[22] S.L. Berger, B. Pina, N. Silverman, G.A. Marcus, J. Agapite, J.L. Regier, S.J. Triezenberg, L. Guarente, Genetic isolation of ADA2: a potential transcriptional adaptor required for function of certain acidic activation domains, Cell 70(2) (1992) 251–65.

[23] A. Saleh, V. Lang, R. Cook, C.J. Brandl, Identification of native complexes containing the yeast coactivator/repressor proteins NGG1/ADA3 and ADA2, J Biol Chem 272(9) (1997) 5571–8.

[24] C. Bian, C. Xu, J. Ruan, K.K. Lee, T.L. Burke, W. Tempel, D. Barsyte, J. Li, M. Wu, B.O. Zhou, B.E. Fleharty, A. Paulson, A. Allali-Hassani, J.Q. Zhou, G. Mer, P.A. Grant, J.L. Workman, J. Zang, J. Min, Sgf29 binds histone H3K4me2/3 and is required for SAGA complex recruitment and histone H3 acetylation, EMBO J 30(14) (2011) 2829–42.

[25] S.L. Sanders, J. Jennings, A. Canutescu, A.J. Link, P.A. Weil, Proteomics of the eukaryotic transcription machinery: identification of proteins associated with components of yeast TFIID by multidimensional mass spectrometry, Mol Cell Biol 22(13) (2002) 4723–38.

[26] P.A. Grant, L. Duggan, J. Cote, S.M. Roberts, J.E. Brownell, R. Candau, R. Ohba, T. Owen-Hughes, C.D. Allis, F. Winston, S.L. Berger, J.L. Workman, Yeast Gcn5 functions in two multisubunit complexes to acetylate nucleosomal histones: characterization of an Ada complex and the SAGA (Spt/Ada) complex, Genes Dev 11(13) (1997) 1640–50.

[27] A. Shukla, S. Lahudkar, G. Durairaj, S.R. Bhaumik, Sgf29p facilitates the recruitment of TATA box binding protein but does not alter SAGA’s global structural integrity in vivo, Biochemistry 51(2) (2012) 706–14.

[28] M. Vermeulen, H.C. Eberl, F. Matarese, H. Marks, S. Denissov, F. Butter, K.K. Lee, J.V. Olsen, A.A. Hyman, H.G. Stunnenberg, M. Mann, Quantitative interaction proteomics and genome-wide profiling of epigenetic histone marks and their readers, Cell 142(6) (2010) 967–80.

[29] A.E. Ringel, A.M. Cieniewicz, S.D. Taverna, C. Wolberger, Nucleosome competition reveals processive acetylation by the SAGA HAT module, Proc Natl Acad Sci U S A 112(40) (2015) E5461–70.

[30] L.A. Boyer, M.R. Langer, K.A. Crowley, S. Tan, J.M. Denu, C.L. Peterson, Essential role for the SANT domain in the functioning of multiple chromatin remodeling enzymes, Mol Cell 10(4) (2002) 935–42.

[31] D.E. Sterner, X. Wang, M.H. Bloom, G.M. Simon, S.L. Berger, The SANT domain of Ada2 is required for normal acetylation of histones by the yeast SAGA complex, J Biol Chem 277(10) (2002) 8178–86.

[32] J. Sun, M. Paduch, S.A. Kim, R.M. Kramer, A.F. Barrios, V. Lu, J. Luke, S. Usatyuk, A.A. Kossiakoff, S. Tan, Structural basis for activation of SAGA histone acetyltransferase Gcn5 by partner subunit Ada2, Proc Natl Acad Sci U S A 115(40) (2018) 10010–10015.

[33] L. Aravind, L.M. Iyer, The SWIRM domain: a conserved module found in chromosomal proteins points to novel chromatin-modifying activities, Genome Biol 3(8) (2002) RESEARCH0039.

[34] C. Qian, Q. Zhang, S. Li, L. Zeng, M.J. Walsh, M.M. Zhou, Structure and chromosomal DNA binding of the SWIRM domain, Nat Struct Mol Biol 12(12) (2005) 1078–85.

[35] N. Tochio, T. Umehara, S. Koshiba, M. Inoue, T. Yabuki, M. Aoki, E. Seki, S. Watanabe, Y. Tomo, M. Hanada, M. Ikari, M. Sato, T. Terada, T. Nagase, O. Ohara, M. Shirouzu, A. Tanaka, T. Kigawa, S. Yokoyama, Solution structure of the SWIRM domain of human histone demethylase LSD1, Structure 14(3) (2006) 457–68.

[36] G. Da, J. Lenkart, K. Zhao, R. Shiekhattar, B.R. Cairns, R. Marmorstein, Structure and function of the SWIRM domain, a conserved protein module found in chromatin regulatory complexes, Proc Natl Acad Sci U S A 103(7) (2006) 2057–62.

[37] Y. Han, J. Luo, J. Ranish, S. Hahn, Architecture of the Saccharomyces cerevisiae SAGA transcription coactivator complex, EMBO J 33(21) (2014) 2534–46.

[38] B.S. Atanassov, Y.A. Evrard, A.S. Multani, Z. Zhang, L. Tora, D. Devys, S. Chang, S.Y. Dent, Gcn5 and SAGA regulate shelterin protein turnover and telomere maintenance, Mol Cell 35(3) (2009) 352–64.

[39] T.L. Burke, J.L. Miller, P.A. Grant, Direct inhibition of Gcn5 protein catalytic activity by polyglutamine-expanded ataxin-7, J Biol Chem 288(47) (2013) 34266–34275.

[40] A. Barrios, W. Selleck, B. Hnatkovich, R. Kramer, D. Sermwittayawong, S. Tan, Expression and purification of recombinant yeast Ada2/Ada3/Gcn5 and Piccolo NuA4 histone acetyltransferase complexes, Methods 41(3) (2007) 271–7.

[41] P.T. Lowary, J. Widom, New DNA sequence rules for high affinity binding to histone octamer and sequence-directed nucleosome positioning, J Mol Biol 276(1) (1998) 19–42.

[42] K. Luger, T.J. Rechsteiner, T.J. Richmond, Expression and purification of recombinant histones and nucleosome reconstitution, Methods Mol Biol 119 (1999) 1–16.

[43] C.E. Berndsen, J.M. Denu, Assays for mechanistic investigations of protein/histone acetyltransferases, Methods 36(4) (2005) 321–31.

[44] C.S. Hughes, S. Moggridge, T. Müller, P.H. Sorensen, G.B. Morin, J. Krijgsveld, Single-pot, solid-phase-enhanced sample preparation for proteomics experiments, Nat Protoc 14(1) (2019) 68–85.

[45] R. Evans, M. O’Neill, A. Pritzel, N. Antropova, A. Senior, T. Green, A. Žídek, R. Bates, S. Blackwell, J. Yim, O. Ronneberger, S. Bodenstein, M. Zielinski, A. Bridgland, A. Potapenko, A. Cowie, K. Tunyasuvunakool, R. Jain, E. Clancy, P. Kohli, J. Jumper, D. Hassabis, Protein complex prediction with AlphaFold-Multimer, bioRxiv (2022) 2021.10.04.463034.

[46] M. Mirdita, K. Schutze, Y. Moriwaki, L. Heo, S. Ovchinnikov, M. Steinegger, ColabFold: making protein folding accessible to all, Nat Methods 19(6) (2022) 679–682.

[47] V. Hornak, R. Abel, A. Okur, B. Strockbine, A. Roitberg, C. Simmerling, Comparison of multiple Amber force fields and development of improved protein backbone parameters, Proteins 65(3) (2006) 712–25.

[48] A. Eberharter, D.E. Sterner, D. Schieltz, A. Hassan, J.R. Yates, 3rd, S.L. Berger, J.L. Workman, The ADA complex is a distinct histone acetyltransferase complex in Saccharomyces cerevisiae, Mol Cell Biol 19(10) (1999) 6621–31.

[49] G. Liu, X. Zheng, H. Guan, Y. Cao, H. Qu, J. Kang, X. Ren, J. Lei, M.Q. Dong, X. Li, H. Li, Architecture of Saccharomyces cerevisiae SAGA complex, Cell Discov 5 (2019) 25.

[50] D. Vasyliuk, J. Felt, E.D. Zhong, B. Berger, J.H. Davis, C.K. Yip, Conformational landscape of the yeast SAGA complex as revealed by cryo-EM, Sci Rep 12(1) (2022) 12306.

[51] P.A. Grant, A. Eberharter, S. John, R.G. Cook, B.M. Turner, J.L. Workman, Expanded lysine acetylation specificity of Gcn5 in native complexes, J Biol Chem 274(9) (1999) 5895–900.

[52] Y.M. Kuo, A.J. Andrews, Quantitating the specificity and selectivity of Gcn5-mediated acetylation of histone H3, PLoS One 8(2) (2013) e54896.

[53] V. Mariani, M. Biasini, A. Barbato, T. Schwede, lDDT: a local superposition-free score for comparing protein structures and models using distance difference tests, Bioinformatics 29(21) (2013) 2722–8.

[54] D. Setiaputra, J.D. Ross, S. Lu, D.T. Cheng, M.Q. Dong, C.K. Yip, Conformational flexibility and subunit arrangement of the modular yeast Spt-Ada-Gcn5 acetyltransferase complex, J Biol Chem 290(16) (2015) 10057–70.

