## Supplemental figures for "The SAGA HAT module is tethered by its SWIRM domain and modulates activity of the SAGA DUB module"

**Supplementary Information**

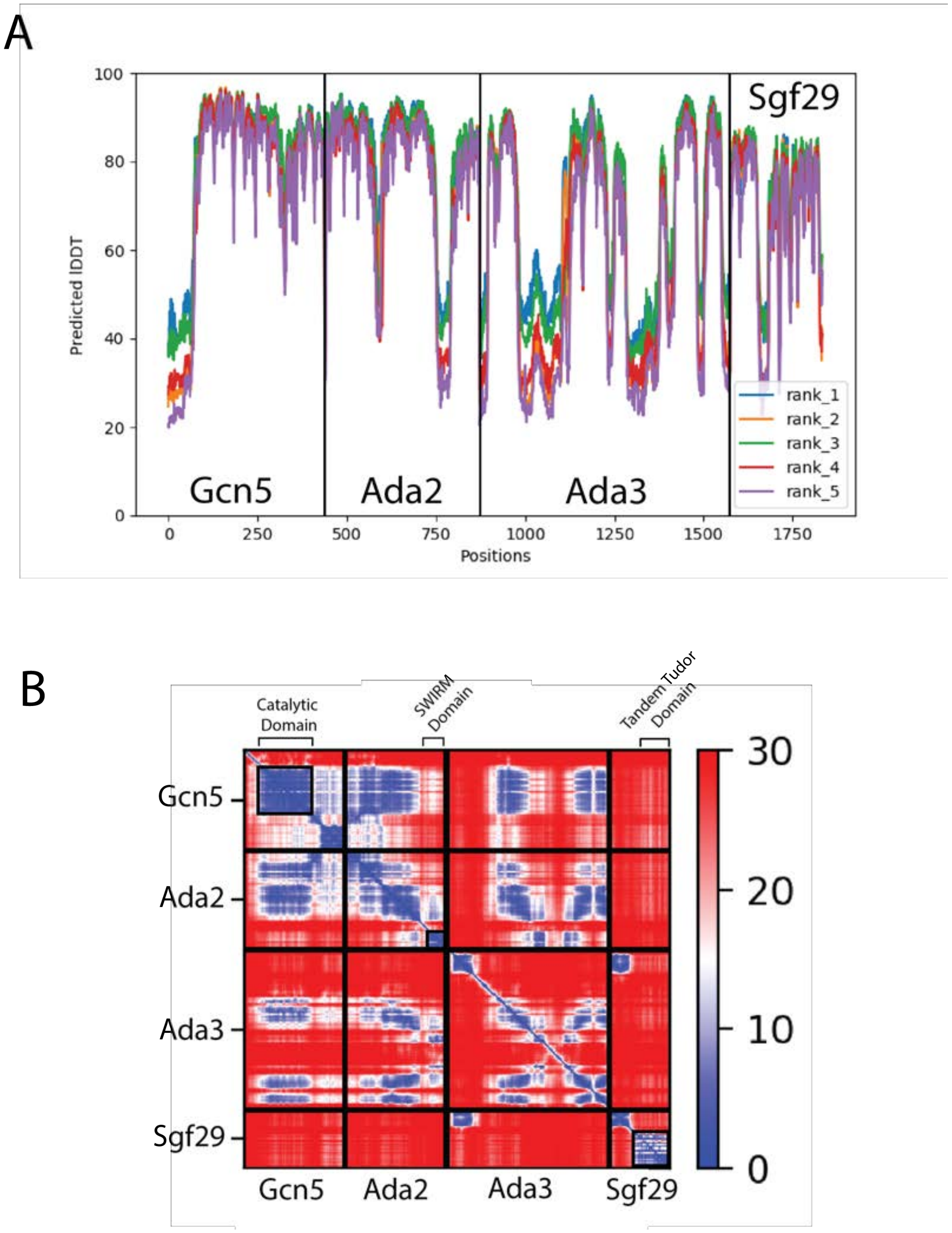


**Supplementary Figure S1.** AF2-Multimer model accuracy metrics. **A)** Predicted Local Distance Difference Test (pLDDT) score of the top five predicted structures of the HAT module. **B)** Predicted Alignment Error (PAE) score of the highest rank model. Areas corresponding to the SWIRM, tandem Tudor, and the catalytic domain are marked.

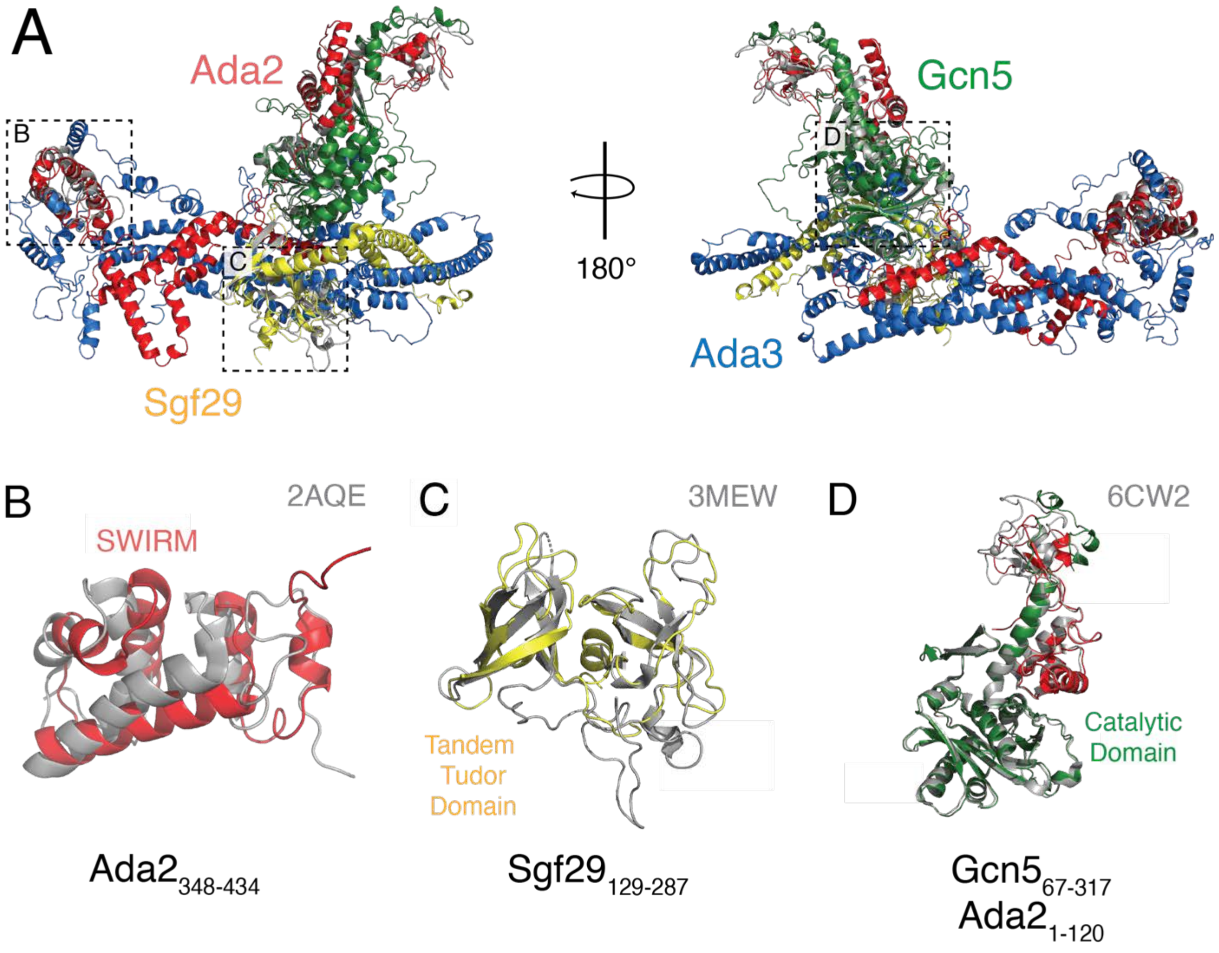


**Supplementary Figure S2.** Comparison of AF2-Multimer model of yeast HAT module against experimentally determined structures of HAT module components. (A) Overview of HAT module with crystal structures superimposed in gray. (B) Structure alignment of the solution structure of Ada2's SWIRM domain (2AQE, gray) and the AF2-Multimer model (red). (C) Structure alignment of the crystal structure of Sgf29's tandem Tudor domain (3MEW, gray) and the AF2-Multimer model (yellow). (D) Structure alignment of the crystal structure of $Ada2_{1\text{-}120}$ and $Gcn5_{67\text{-}317}$ (6CW2, gray) and the AF2-Multimer model (green and red).
